# Detection of Dugbe virus from ticks in Ghana

**DOI:** 10.1101/2022.04.04.486928

**Authors:** Charlotte Adwoa Addae, Michael Wiley, Catherine Pratt, Jeffrey W. Koehler, Seth Offei Addo, Mba Mosore, Clara Yeboah, Bright Agbodzi, Danielle Ladzekpo, Janice A. Tagoe, Eric Behene, Isaac Adrah, Courage Defeamekpor, Osbourne Quaye, Andrew Letizia, Shirley Nimo-Paintsil, Hanayo Arimoto, Joseph W. Diclaro, Samuel Dadzie

## Abstract

Vector-borne pathogens historically have impacted U.S. warfighters and active-duty personnel stationed both domestically and globally during deployments or forward operations. Tick-borne diseases continue to spread to new geographical regions affecting both animal and human health. In Ghana, there is limited information on the circulating tick-borne pathogens and their risk of infections. This military-to-military vector surveillance study focused on seven sites in Ghana: Navrongo, Airforce Base, 6th Battalion Infantry, Air Borne Force, Army Recruit Training School, 1st Battalion Infantry and 5th Battalion Infantry for detection of tick-borne pathogens. Ticks from these sites were collected by hand-picking with a pair of forceps from domesticated animals including cattle, sheep, goats and dogs. A total of 2,016 ticks were collected from two main ecological zones; the northern Sahel savannah and the coastal savannah. *Amblyomma variegatum* was the predominant species, accounting for 59.5% of the collected ticks. Next-generation sequencing allowed for Dugbe virus whole genome detection. This study reports the second detection of Dugbe virus in Ghana, which is closely related to Dugbe virus strains from Kenya and Nigeria. This study (sponsored by the Armed Forces Health Surveillance Branch-Global Emerg-ing Infections Surveillance Section) aimed to better inform Force Health Protection (FHP) decisions within the U.S. AFRICOM area of responsibility, the Ghana Armed Forces, and enhance global health security countermeasures. Further surveillance needs to be conducted within the country to determine the distribution of tick-borne pathogens to formulate effective control measures.

**Author summary:** The prevalence of Dugbe virus in tick species within Ghana was investigated. This study involved seven sampling sites that covered two ecological zones, the northern Sahel savannah and the coastal savannah. About 2,000 ticks were collected from cattle, sheep, goats and dogs and identified using a dissecting microscope. The most predominant tick species was *Amblyomma variegatum* (59.5%) also known as the tropical bont tick. Using Next-generation sequencing, the full genome of Dugbe virus was for the first time in Ghana identified in *A. variegatum*. This positive *Amblyomma variegatum* was collected from the Greater Accra region of Ghana within a military site. The virus was found to be related to Dugbe virus strains from Kenya and Nigeria. Findings from this study show that the trade of livestock across borders facilitates the spread of tick-borne pathogens hence there is a need to enforce measures that prevent the importation of potentially harmful ticks and tick-borne pathogens.

## 1. Introduction

Tick-borne diseases (TBDs) represent a significant threat to human health, especially in regions where close contacts of people with livestock occur. Previous studies in Ghana have found evidence of multiple TBDs including diseases that are caused by Crimean Congo haemorrhagic fever virus (CCHFV) [1], Dugbe virus [2], Rickettsia [3], and Coxiella [4,5] [4]. However, the prevalence and transmission risk of these pathogens in Ghana are not well understood.

Ghana’s location on the West African coast makes it vital in U.S. AFRICOM’s effort in its engagement with West African partners. The collaboration between the U.S. Naval Medical Research Unit - No.3 (NAMRU-3), Noguchi Memorial Institute for Medical Research (NMIMR), and the Navrongo Health Research Centre (NHRC) supports the U.S. military in its effort to combat multiple priority pathogens, such as TBDs, that potentially cause diseases among U.S. and Ghanaian soldiers deployed in the austere environments.

Dugbe virus (DUGV) is a single-stranded, negative-sense RNA virus that belongs to the family Nairoviridae, genus Orthonairovirus [6], and the Nairobi sheep disease virus serogroup [7,8]. Dugbe virus has been found throughout Africa including Kenya [9,10], Nigeria [11], and Ghana [2] and is primarily transmitted by the ixodid tick Amblyomma variegatum [12,13], but can also be spread by Rhipicephalus and Haemaphysalis tick species [14]. Human infection with DUGV is usually less severe, causing a mild flu-like disease [14,15].

This study is part of a larger project aimed to collect, enumerate, and morphologically identify ticks from livestock located in two main ecological zones in Ghana and characterize potential pathogens that are present in the ticks.

## 2. Materials and Methods

### 2.1 Study Sites and Tick Collection

Two main ecological zones (the northern Sahel savannah and the coastal savannah) in Ghana were surveyed in this study (Fig 1). The study protocol was reviewed and approved by the Institutional Review Board of the Noguchi Memorial Institute for Medical Research (NMIMR) (Decision number: 110/15-16). Informed verbal consent was obtained from the owners, herdsmen, and farm managers of the livestock farms before sample collection. Ticks were collected from the Sahel savannah in August 2017 and March 2018, and monthly from December 2017 to March 2018 in the coastal savannah.

**Fig 1:**
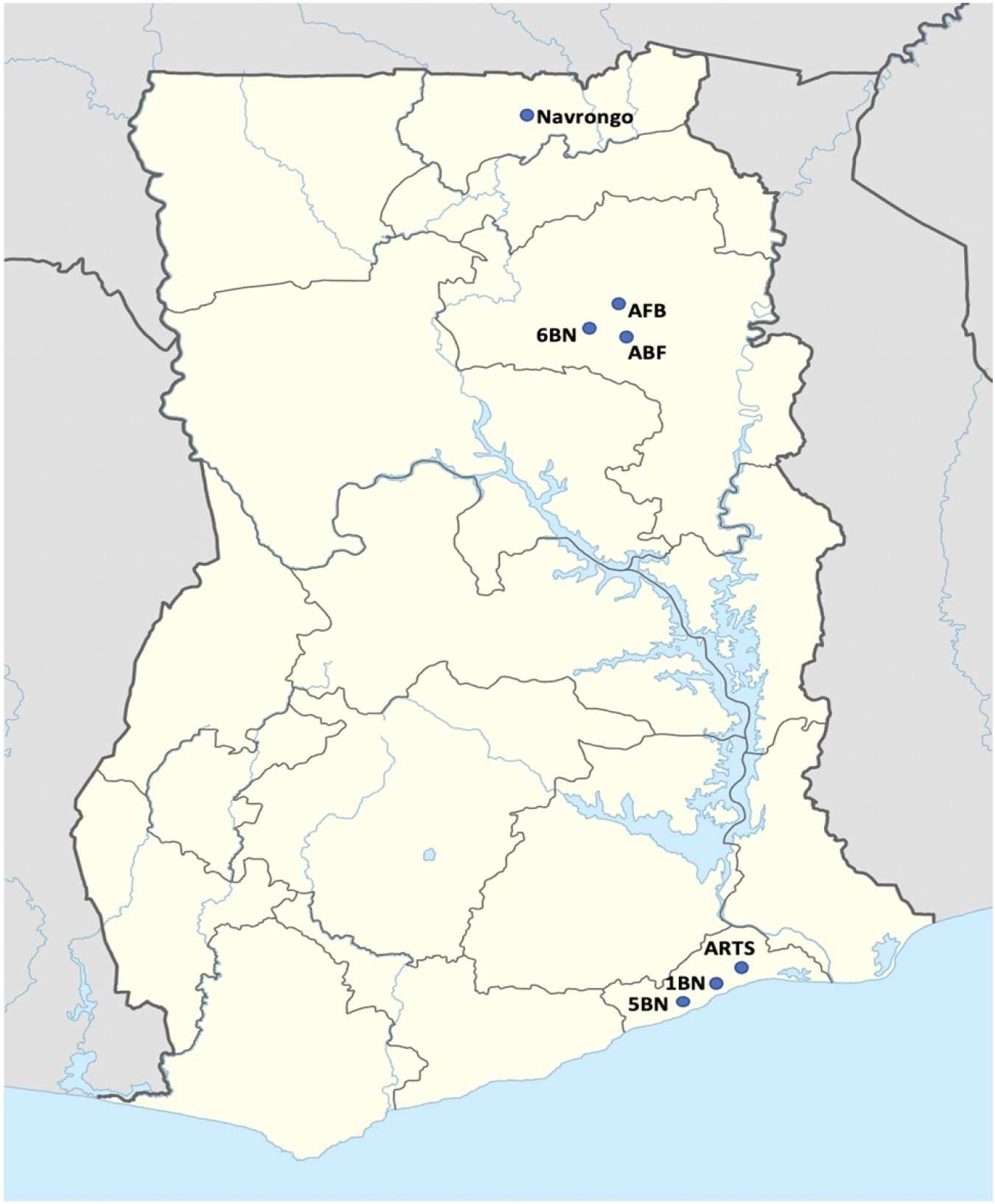
Map of Ghana showing the various study sites; Navrongo, Air Force Base (AFB), 6th Battalion Infantry (6 BN), Air Borne Force (ABF), Army Recruit Training School (ARTS), 1st Bat-talion Infantry (1 BN) and 5th Battalion Infantry (5 BN).[16].

### 2.2 Sample processing

Ticks collected from cattle, sheep, goats and dogs were placed in RNAlater (ThermoFisher Scientific, Waltham, USA) and transported to NMIMR for further analysis. Morphological identification was performed using the African Ixodidae identification keys [17]. Identified ticks were pooled, based on species, gender, study site, and animal source, into either three males or two females. The pooled samples were homogenized in a Mini-Beadbeater-96 (BioSpec, Bartlesville, OK, USA) using specific beads that were 0.1 mm and 2.0 mm in diameters and lysis buffer as previously described [18].

### 2.3 Viral detection using Next-generation Sequencing

Total nucleic acid was extracted from the homogenized samples using QIAamp Viral RNA Mini Kit (QIAGEN, Valencia, CA, USA) by following the manufacturer’s instructions [19]. Libraries were prepared from the extracted nucleic acid using the TruSeq RNA Access Library Prep Kit (Illumina, San Diego, CA, USA). Wide range pan viral probes were used to target unknown RNA viral genomes and detect virus types. The libraries were amplified for 17 cycles and the products were quantified by qPCR using the KAPA syber fast universal qPCR kit (KAPA Biosystems, KK4824) [20] on an Agilent 2100 bioanalyzer (Agilent Technologies, Santa Clara, USA). The libraries were pooled and sequenced on a MiSeq sequencing platform (Illumina, San Diego, CA, USA) at 2 × 151-bp paired-end sequencing with v3 chemistry.

### 2.4 Phylogenetic analysis

The sequence data were assembled using BLAST reference sequences in the Nairovirus family and other virus sequences from different serogroups in the Orthonairovirus genus that were obtained from GenBank were aligned with the assembled genes (sequenced data). Alignments of the DUGV L, M and S segments were done separately using muscle in MEGA7 [21]. For the L segment alignment, the DUGV L segment sequence that was detected in Pokuase by Kobayashi and his colleagues [2] was added to the analysis. The resulting alignments were analysed using a maximum likelihood algorithm and phylogenetic trees were `created using MEGA7 with 1000 bootstrap replications.

Phylogenetic trees for the three viral sequences were rooted with Bunyamwera virus of the Nairovirus family and Bunyamwera serogroup [22] because the virus is considered a prototype for other viruses in the family [23].

## 3. Results

### 3.1 Tick collection

A total of 2016 ticks were collected from both ecological areas with 82% of the total number of ticks collected from cattle. Approximately, 60% of all the ticks collected were Amblyomma variegatum, with the rest being Rhipicephalus and Hyalomma species. Male and female ticks accounted for 65% and 35%, respectively, of the total number of ticks collected (Table 1).

**Table 1:** Distribution of tick species, sex, and animal source from the Northern Sahel and Coastal savanna ecological zones.

| Tick species | Animal source | Northern Sahel savannah |  | Coastal savannah |  | Total |
| --- | --- | --- | --- | --- | --- | --- |
|  |  | Male | Female | Male | Female |  |
| *Amblyomma variegatum* | Cattle | 677 | 72 | 303 | 146 | 1198 |
|  | Dog | 2 | 0 | 0 | 0 | 2 |
| Sub Total |  | 679 | 72 | 303 | 146 | 1200 |
| *Hyalomma rufipes* | Cattle | 8 | 4 | 20 | 20 | 52 |
| *Hyalomma truncatum* | Cattle | 86 | 40 | 1 | 0 | 127 |
| *Rhipicephalus sanguineus* | Cattle | 25 | 29 | 4 | 37 | 95 |
|  | Dog | 167 | 145 | 0 | 0 | 312 |
|  | Goat | 0 | 6 | 0 | 0 | 6 |
| Sub Total |  | 192 | 180 | 4 | 37 | 413 |
| *Rhipicephalus evertsi* | Cattle | 2 | 0 | 0 | 7 | 9 |
|  | Sheep | 8 | 2 | 0 | 0 | 10 |
|  | Goat | 1 | 0 | 0 | 0 | 1 |
| Sub Total |  | 11 | 2 | 0 | 7 | 20 |
| *Rhipicephalus spp.* | Cattle | 0 | 0 | 0 | 189 | 189 |
|  | Dog | 0 | 0 | 10 | 4 | 14 |
| Sub Total |  | 0 | 0 | 10 | 193 | 203 |
| *Rhipicephalus boophilus* | Cattle | 0 | 0 | 1 | 0 | 1 |
| Total |  | 976 | 298 | 339 | 403 | 2016 |

### 3.2 Virus detection and characterization

Using next-generation sequencing, a full genome sequence for DUGV was found in the *A. variegatum* pool from the Michel Camp in the Greater Accra Region. Phylogenetic analysis revealed that the identified DUGV strain was closely related to DUGV strains from Kenya and Nigeria (Figs 2A, 2B and 2C). The L sequence of the virus obtained in this study reflects a close relationship (99%) with the Dugbe strain (L segment, LC193450) that was detected in Ghana [2]. The M segment was also closely related by 92% and 73% to the Kenya and Nigeria strains. The S segment of the viral genome had a significant closeness to the Kenyan, Nigerian and Senegal strains (98%, 99% and 60% respectively) as compared to the M and L segment which were much similar to other strains (Middle Eastern and Asian strains. The S segment was also closely related to the Kenya and Nigeria strains.

**Fig 2A:**
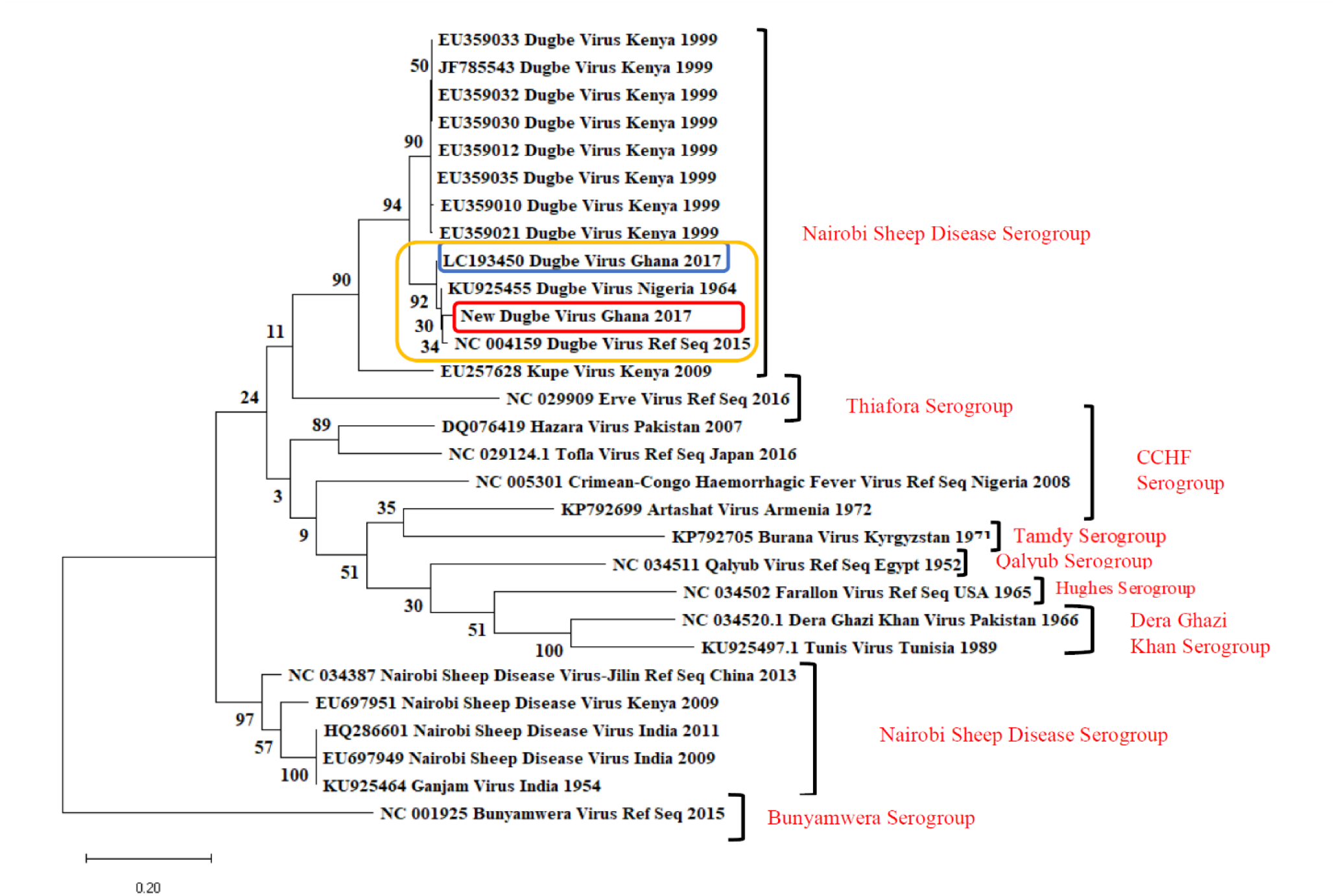
Phylogenetic analysis of gene segments of DUGV detected in Ghana. The phylogenetic trees are for the (A) L segment, (B) M segment, and (C) S segment. DUGV strains and reference strains of other viruses in the Nairoviridae family were used for the analysis. The DUGV that was detected in this study is shown in the red rectangle. The first virus detected in Pokuase is shown in blue. Bunyamwera virus is the root of the tree.

**Fig 2B:**
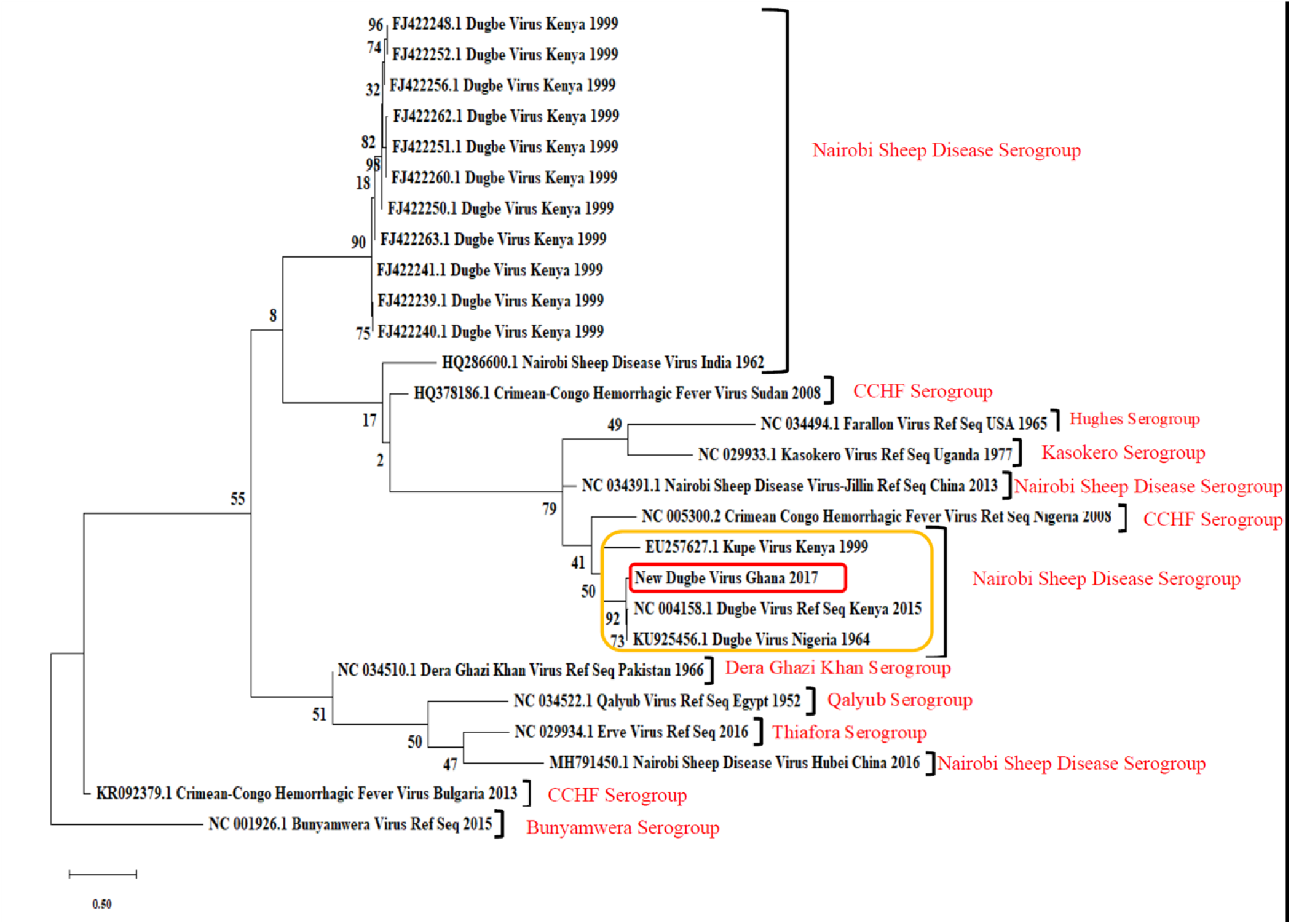
Phylogenetic tree showing the relationship between the M segment of the new DUGV detected in Ghana, other DUGV strains and reference strains of other viruses in the Nairoviridae family. The M segment is closely related to the Dugbe virus strain from Nigeria with accession number KU925456.1, and the reference Dugbe virus strain (NC004158.1) as shown with the yellow rectangle. The new DUGV detected is shown in the red rectangle. Bunyamwera virus is the root of the tree.

**Fig 2C:**
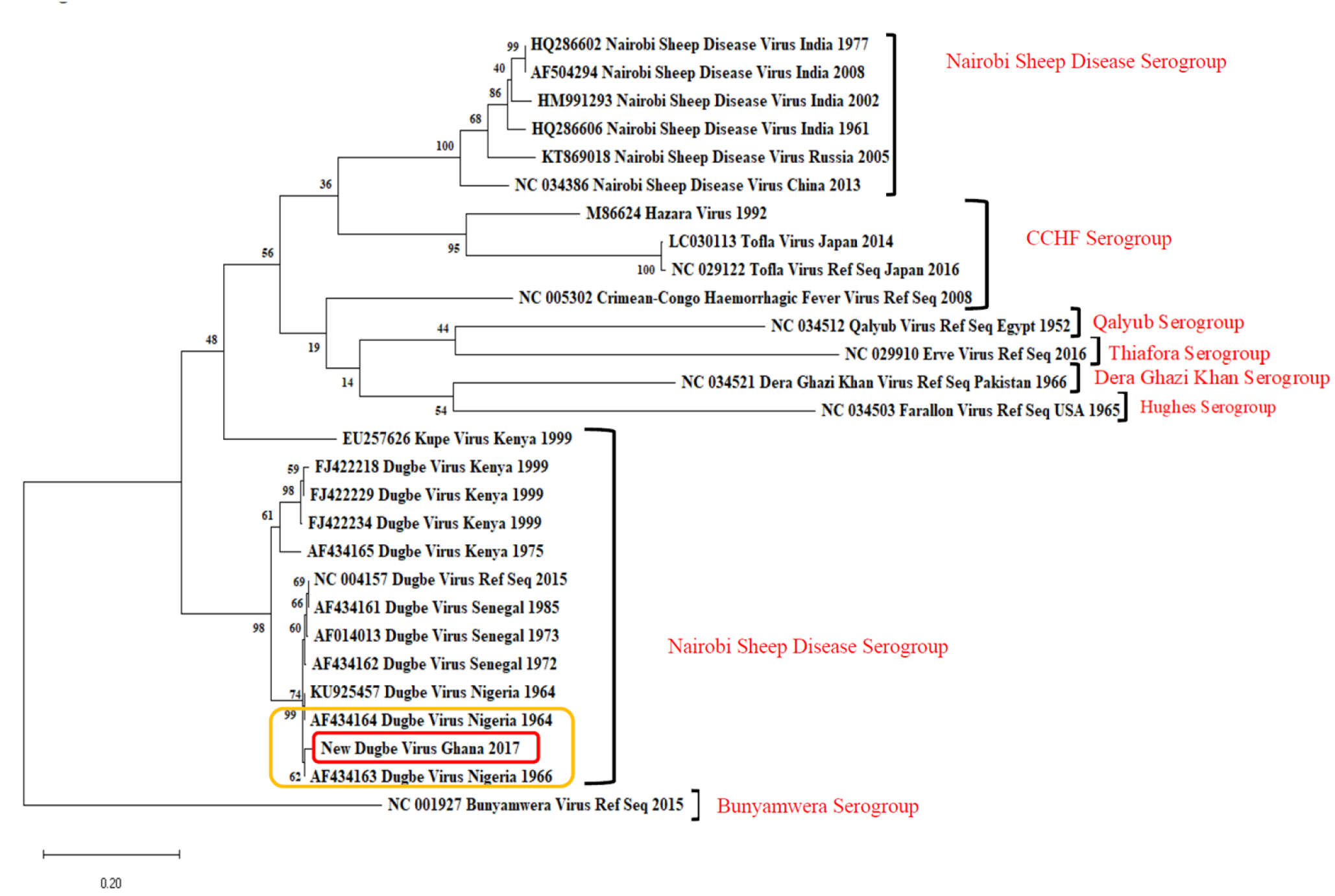
Phylogenetic tree showing the relationship between the S segment of the new DUGV detected in Ghana, other DUGV strains and reference strains of other viruses in the Nairoviridae family. The S segment is closer to the Dugbe strain from Nigeria, AF434163, and is in the same clade with DUGV strains from Senegal, Kenya, and Nigeria. The new DUGV detected is shown in the rectangle. Bunyamwera virus is the root of the tree.

## 4. Discussion

In this study, 3 genera of ticks including Amblyomma, Rhipicephalus and Hyalomma were identified from two ecological zones, with Amblyomma variegatum as the most predominant tick species in both ecological zones. In Africa, A. variegatum is the common and most widely distributed tick that affects livestock including cattle, goats, and sheep [17]. This particular species occurs in a wide range of climates and transmits pathogens that cause significant economic loss in ruminant production and infections in humans [24,25]. Previous studies from Ghana have reported the wide distribution of A. variegatum on ruminants [26-28] and are consistent with the trend observed in this study. It was also observed that Rhipicephalus sanguineus occurred mostly on dogs, which have been established as the primary hosts, although this tick species can also be found on other animals and humans [29,30].

DUGV is a pathogenic Nairovirus that is endemic in arid regions, one of the frequently occurring tick-borne viruses in Africa, and has been associated with tick infestation in livestock [31,32]. Dugbe virus has been detected in ticks from an abattoir in Nairobi [10] and more recently in tick species sampled in Ghana [2]. In pregnant livestock, stillbirths may occur when infected with the virus causing economic loss [7]. Although DUGV is predominantly found in animals, human infections can occur through the bite of an infected tick or contact with the infected bodily fluids of an animal or human [33]. Nairo-viruses are considered one of the rapidly emerging tick-borne viruses globally and in Africa [34]. Although the viruses have been linked to diseases in livestock, the infection has been isolated from a person with mild febrile disease [35]. Domestic animals are known to maintain the viral cycle and therefore behavior that increases the risk of exposure through bites and handling of carcasses by humans, especially animal handlers should be discouraged.

Sequencing analysis of the L, M and S segments of the DUGV strain from this study showed close relatedness to other strains that have been detected in both Ghana and Nigeria [36]. The virus has been frequently isolated from tick infecting market livestock [37], and the trade of livestock across African countries [36], pathogens can migrate from endemic regions to new locations. It is important to assess the effects of the characterized pathogen on livestock health, as a potential outbreak could lead to economic loss at a national level. Ticks across the country could also harbour other pathogens that could be of public health importance, and with the recent re-emergence of tick-borne viruses in Africa, surveillance efforts targeting major livestock communities within the country will be essential. In the meanwhile, control efforts such as the use of acaricides to reduce the tick population need to be employed to prevent the spread of infections.

## 5. Conclusions

The findings from this study indicate the presence of diverse tick species across the sampling sites with A. variegatum as the predominant species and report the second detection of Dugbe virus in infected ticks in Ghana. This is the first report of the full genome sequence for Dugbe virus in Ghana. Data from this study will be used to provide accurate reporting across the GEIS surveillance network and to inform FHP decision making in both U.S. and Ghana Armed Forces and guide Ghana’s public health infectious diseases policy. It is important to conduct further studies in the country to determine the distribution of tick-borne pathogens and assess the risk of transmission to humans, especially animal handlers and livestock owners.

## Author Contributions

***Charlotte Adwoa Addae***: Methodology, investigation, writing—original draft preparation, writing— review and editing.

***Michael Wiley***: investigation, data curation, writing—review and editing.

***Catherine Pratt***: investigation, data curation, writing—review and editing.

***Jeffrey W. Koehler***: investigation, data curation, writing—review and editing.

***Seth Offei Addo***: Methodology, investigation, writing—original draft preparation, writing—review and editing.

***Mba Mosore***: Methodology, investigation, writing—original draft preparation, writing—review and editing.

***Clara Yeboah***: Methodology, investigation, writing—review and editing.

***Bright Agbodzi***: Methodology, investigation, writing—review and editing.

***Danielle Ladzekpo***: investigation, writing—review and editing.

***Janice A. Tagoe***: Methodology, investigation, writing—review and editing.

***Eric Behene***: data curation, writing—review and editing.

***Isaac Adrah***: Methodology, investigation, writing—review and editing.

***Courage Defeamekpor***: Methodology, investigation, writing—review and editing.

***Osbourne Quaye***: investigation, writing—review and editing, supervision.

***Andrew Letizia***: writing—review and editing, supervision, project administration, funding acquisition.

***Shirley Nimo-Paintsil***: Conceptualization, Methodology, writing—review and editing, project administration.

***Hanayo Arimoto***: writing—review and editing, supervision.

***Joseph W. Diclaro II***: Conceptualization, writing—review and editing, supervision, project administration, funding acquisition.

***Samuel Dadzie***: writing—review and editing, supervision, project administration

## Funding/Disclaimer

CDR Andrew Letizia, LCDR Joseph Diclaro, and LT Hanayo Arimoto are military Service members, and Dr Shirley Nimo-Paintsil is an employee of the U.S. Government. This work was prepared as part of their official duties. Title 17, U.S.C. §105 provides that copyright protection under this title is not available for any work of the U.S. Government. The views expressed in this article are those of the authors and do not necessarily reflect the official policy or position of the Department of the Navy, Department of Defense, nor the U.S. Government. This work was supported by the Armed Forces Health Surveillance Branch and its Global Emerging Infections Surveillance and Response (GEIS) Section (funding year 2017, ProMIS ID P2031_16_N3). The study protocol was reviewed and approved by the Institutional Review Board of the Noguchi Memorial Institute for Medical Research (NMIMR) (Decision number: 110/15-16). Informed verbal consent was obtained from the owners, herdsmen, and farm managers of the livestock farms before sample collection.

## Data Availability Statement

This article includes all of the data that supports the findings.

## Acknowledgements

Special thank you to the staff of the Veterinary Department of the Ghana Armed forces, 37 Military Hospital, Accra for their support in tick collection and assistance with the safe handling of the animals from which ticks were collected for this study. The authors are also grateful to the NMIMR, NECE, NAMRU-3 Ghana Detachment and Navrongo Health Research Center Entomology teams for their support and collaboration. We are also indebted to the USAMRIID team for their assistance in reverse transcription real-time PCR assays and sequencing technical expertise.

